# Comparative Analysis of FOXP2 Expression in the Thalamus of Mice, Rats, and Macaques: Implications for the Evolution of Language Circuits

**DOI:** 10.1101/2025.11.23.689322

**Authors:** Blanca Sánchez-Moreno, Alicia Uceda-Heras, María de la Fuente-Fernández, Miguel Ángel García-Cabezas, Carmen Cavada, Javier Gilabert-Juan

## Abstract

FOXP2 is a transcription factor essential for the development and function of neural circuits involved in language. Although its expression has been extensively characterized in the cortex and basal ganglia, its organization within the adult thalamus remains poorly understood. In this study, we present a comparative analysis of FoxP2 protein expression across thalamic nuclei in mice, rats, and macaques, with a focus on nuclei associated with higher-order cognitive functions and language-related circuits in humans. We found that FoxP2 is expressed in most thalamic nuclei across species, with a consistent absence in the reticular nucleus and zona incerta. Expression was highest in midline and intralaminar nuclei, whereas the anterior group showed low and variable expression among species. Macaques exhibited broader and, in some nuclei, more intense FoxP2 expression, particularly in associative regions such as the pulvinar, lateral geniculate, and parts of the ventral group, indicating increased specialization of thalamocortical pathways. This distribution suggests a conserved role for FoxP2 in shaping thalamic circuits supporting sensorimotor integration, attention, memory, and linguistic processing. Phylogenetic comparisons further indicate that enhanced FoxP2 expression in associative thalamic territories in primates, likely intensified in humans, may have contributed to the evolution of neural circuits required for speech and language. These findings provide molecular and anatomical insights into how FoxP2 helps organize thalamocortical networks relevant both to language function and to neuropsychiatric disorders involving thalamocortical dysconnectivity.

## Introduction

FOXP2 is a transcription factor belonging to the forkhead box (FOX) family, characterized by a conserved forkhead DNA-binding domain and additional regions mediating dimerization and transcriptional regulation (Lai et al., 2001; Shu et al., 2001). As a nuclear protein, FOXP2 regulates the expression of a wide range of target genes involved in neuronal differentiation, neurite outgrowth, and synaptic plasticity (den Hoed et al., 2021). Mutations in *FOXP2* were first described in the KE family (Lai et al., 2001), in which affected members exhibited developmental verbal dyspraxia, marked by deficits in speech motor control, language processing, and grammatical skills (Vargha-Khadem et al., 1998). These findings positioned FOXP2 as a key molecular entry point into the genetic and neurobiological foundations of language. FOXP2 is highly expressed in distinct neuronal populations within cortico-striato-thalamo-cortical circuits, suggesting a conserved role in motor sequencing and sensorimotor integration (Fisher & Scharff, 2009; Vargha-Khadem et al., 2005).

The thalamus is a critical hub supporting both linguistic processing and broader cognitive functions (Klostermann et al., 2013). Ischemic or hemorrhagic lesions can result in thalamic aphasia (Fritsch et al., 2022), underscoring its involvement in speech and language. Although the specific contribution of individual thalamic nuclei remains incompletely understood, the ventral anterior (VA), ventrolateral (VL), intralaminar, and pulvinar nuclei have been most consistently implicated in language functions (Barbas et al., 2013; Crosson, 2013, 2021a; Llano, 2013). Other nuclei, such as the reticular, mediodorsal (MD), and midline nuclei, are thought to support language indirectly through their roles in memory, attention, and executive functions (Crosson, 2013; Perea-Bartolomé & Ladera-Fernández, 2004). Together, these findings highlight the thalamus as a dynamic integrator within the distributed neural network that underlies speech and linguistic cognition.

Within this context, FoxP2 expression in the thalamus becomes particularly relevant. FoxP2 is expressed from early embryonic stages (Alhesain et al., 2025; Lai et al., 2003), displaying a conserved spatial and temporal pattern that persists into adulthood with varying intensity across different nuclei (Lai et al., 2003; Takahashi et al., 2008). Functional studies indicate that FoxP2 contributes to thalamic patterning and the specification of regional thalamic identities, guiding the formation and refinement of thalamocortical projections (Alhesain et al., 2025; Ebisu et al., 2017). Through these mechanisms, FoxP2 may influence the development of the very circuits that support word finding, speech sequencing, and the integration of attention and memory during language tasks. Despite these advances, the specific distribution of FoxP2 expression within the mature thalamus remains poorly characterized. Most research has focused on cortical and striatal regions, leaving the molecular organization of thalamic language-related circuits largely unexplored.

In this study, we present a comparative analysis of FoxP2 protein expression in the thalamus of mice, rats, and macaques, focusing on nuclei associated with higher-order cognitive functions, including those linked to language in humans. By examining both the nuclear distribution of FoxP2 and the thalamocortical projection patterns of these nuclei, this work aims to relate the expression profiles of this transcription factor to the functional architecture of thalamic circuits that have evolutionarily contributed to the development of speech and cognition.

## Materials and Methods

### Animals

Tissue from three macaques (two males of *Macaca Mulatta* and one female of *Macaca nemestrina)*, three male rats (*Rattus norvergicus*; Wistar) and three male mice (*Mus musculus*; C57BL/6) was used to study the distribution of FoxP2 in the thalamus. All animals used were adult housed under controlled temperature and a 12-h light/dark cycle with food and water available ad libitum. The ethics protocol under which the experiments were approved was PROEX 209.7/22 and CEI-39-852. All animal experimentation was conducted in accordance with Directive 2010/63/EU of the European Parliament and of the Council of 22 September 2010 on the protection of animals used for scientific purposes and was approved by the Committee on Bioethics of the Universidad Autónoma de Madrid. Every effort was made to minimize the number of animals used and their suffering.

Animals were deeply anesthetized with pentobarbital (Dolethal, Vetoquinol) perfused transcardially with a 4% paraformaldehyde (Panreac) solution in phosphate buffer (PB; 0.1 M, pH 7.4). Brains were then extracted, cryoprotected in a 30% sucrose solution and sectioned coronally at 50 μm using a sliding microtome for rodents and 40 μm coronal sections for macaques. 10 series were collected from rat and macaque brains, and 4 series were collected from mice brains. One series from each was processed for acetylcholinesterase histochemistry, and another for cresyl-violet staining to identify thalamic nuclei.

### Immunohistochemistry

Immunohistochemistry was performed on free-floating macaque brain sections as follows: Sections were washed in TBS 0.1M, followed by two 10-minute washes with endogenous peroxidase inactivation solution (TBS with 3% H2O2 and10% methanol). After further washes, antigen retrieval was performed in a 10 mM sodium citrate buffer (pH 6.0) for 45 minutes at 80°C. Sections were blocked for 4 hours at 4°C with TBS containing 20% normal goat serum (NGS, Biowest), 5% bovine serum albumin (BSA, Sigma-Aldrich), 0.4% Triton X-100, then incubated with the primary antibody mouse anti-FoxP2 (1:500, ThermoFisher, CL5312) in the same buffer containing 0.2% Triton X-100 three overnights at 4°C. The secondary antibody goat anti-mouse (1:200, Merck, AP181B) was applied for 3 hours at room temperature, followed by incubation in avidin–biotin complex for 1 hour. Labeling was visualized with 3,3′-diaminobenzidine (DAB, Sigma-Aldrich) and H₂O₂, and sections were mounted in 0.33 M phosphate buffer and coverslipped with DEPEX (SERVA Electrophoresis GmbH).

For rat and mouse sections, the same protocol was used with the following modifications: PBS with 1% Triton X-100 was used for washes; antigen retrieval was shortened to 15 minutes; blocking solution contained 10% NGS in PBS with 1% Triton X-100; incubation buffer contained 3% NGS in PBS with 1% Triton X-100; primary antibodies were incubated for one overnight, and secondary antibodies for 1 hour.

### Image preparation and thalamic nuclei delimitation

Images of FoxP2 immunostaining in the thalamus were acquired as mosaics of 20x-objective pictures using Neurolucida software (MBF Bioscience, USA) on a personal computer connected to a Zeiss Axioskop microscope (Zeiss, Germany) through a CX9000 camera (MBF Bioscience, USA) and a motorized microscope stage (MBF Bioscience, USA; Heidenhain Corporation, USA). Image resolution was set to 0.5 pixels/μm for rat and mouse sections and 1 pixel/μm for macaque sections

Images of AChE and Nissl stainings were obtained in the same manner, using mosaics of 2.5× objective images. Thalamic nuclei were delineated with reference to The Mouse Brain in Stereotaxic Coordinates (5th Edition) (Paxinos & Franklin, 2019), The Rat Brain in Stereotaxic Coordinates (7th Edition) (Paxinos & Watson, 2013), and The Thalamus of the Macaca Mulatta (Olszewski, 1952) and The Rhesus Monkey Brain in Stereotaxic Coordinates (Paxinos et al., 2000).

### FoxP2+ cell density and intensity analysis

FoxP2 + cell density and intensity were measured using ImageJ. To estimate cell density, the thalamic nuclei were outlined to measure their area and the number of FoxP2+ cells. An average cell density was then calculated for each nucleus. Based on the highest observed cell density, we assigned a density level of 0 to the nuclei with densities less than 5% of this maximum. The remaining range (from 5% to the maximum density) was divided into five equal intervals, and each thalamic nucleus was assigned a density level according to its value.

To assess FoxP2+ cell intensity, between one and three squared sections (5.000 μm2 for mice, or 20.000 μm2 for rats and monkeys) were selected from each thalamic nucleus. These images were processed using a custom ImageJ macro to calculate the mean gray value of the cells and the background. A cell-to-background intensity ratio was calculated to account for differences in background across nuclei. An average intensity ratio was determined for each nucleus. Using the highest and lowest intensity values, we divided the range into five equal levels and assigned an intensity level to each thalamic nucleus accordingly.

## Results

Overall, FoxP2 expression patterns in the thalamus were relatively conversed across the species studied (Fig 1). FoxP2 was expressed across all thalamic nuclei of macaques and in almost all thalamic nuclei of rats and mice, except for the reticular (R) nucleus and the zona incerta (ZI). Expression was highest in the midline and intralaminar nuclei and lower in the anterior group. Below, we describe FoxP2 expression density and intensity across thalamic nuclei. To provide a clearer overview of the results, we present a summary figure depicting FoxP2 cell density and intensity across thalamic nuclei (Fig 2), as well as summary tables indicating FoxP2+ cell density and intensity across all studied thalamic nuclei (Supplementary Tables 1-3).

**Fig 1:**
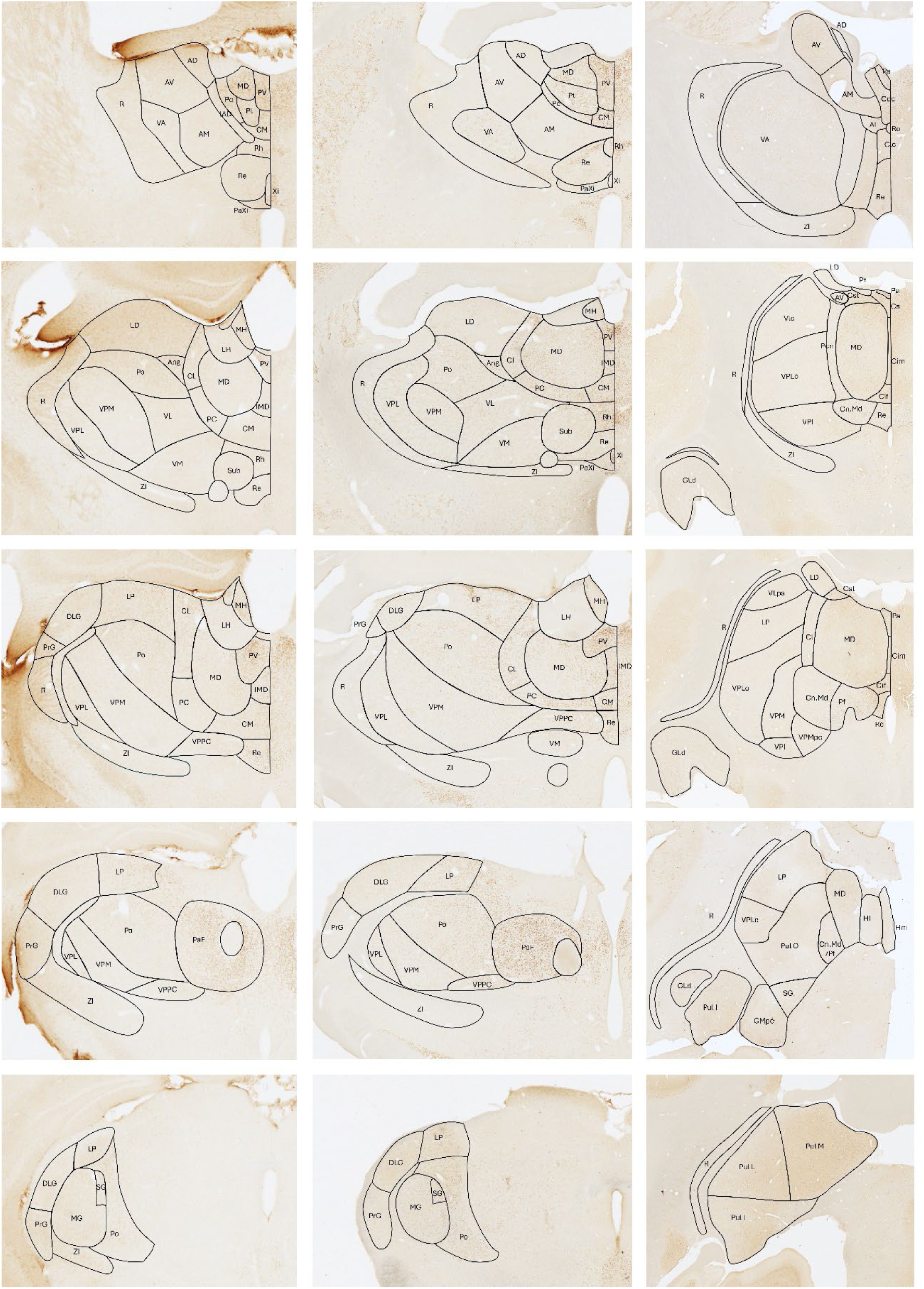
FoxP2 immunohistochemistry in coronal thalamic sections of mice (1), rats (2), and macaques (3). Five representative sections per species, spanning the anterior-posterior axis, are shown. Thalamic nuclei have been delineated. Scale bars: 1.000 μm.

**Fig 2.**
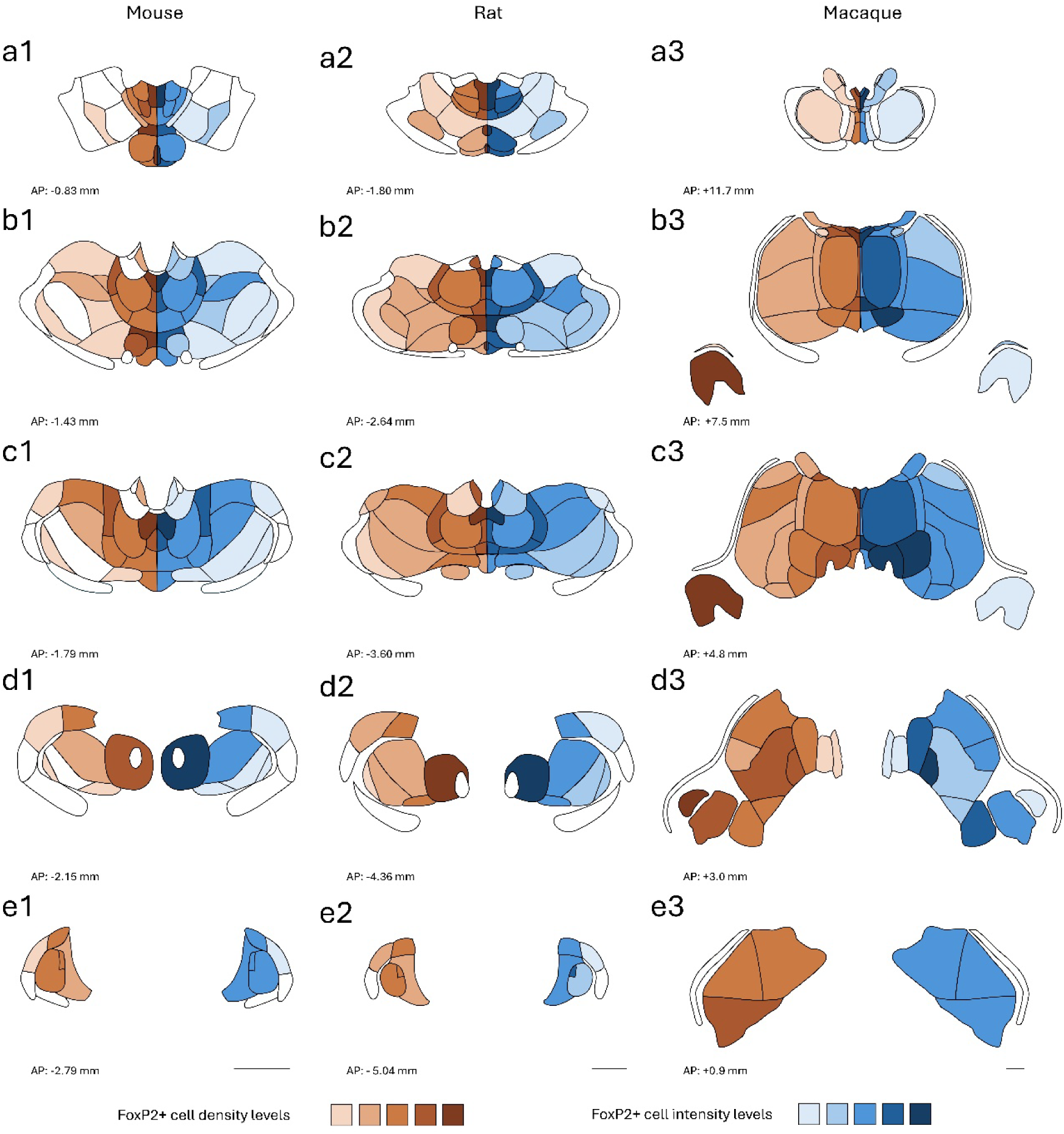
Graphical summary of FoxP2-positive cell density (brown, left) and intensity (blue, right) in coronal thalamic sections of mice (1), rats (2), and macaques (3). Five representative sections per species, spanning the anterior-posterior axis, are shown. Scale bar: 1.000 μm

To enable cross-species comparisons, macaque thalamic nuclei were grouped into the following groups: midline, intralaminar, anterior, ventral and posterior groups, as well as the mediodorsal nucleus, the geniculate nuclei and the pulvinar group (present only in macaques).

### Midline group

Across all species, FoxP2 expression was the highest in the midline group. In mice and rats, the paraventricular (PV), rhomboid (Rh) and xiphoid (Xi) nuclei showed the greatest cell density and labelling intensity. In macaques, the highest expression occurred in the paraventricular (Pa), central superior (Cs) and central inferior (Cif) nuclei. In mice, the lowest expression was observed in the interanteromedial (IAM) nucleus, with medium levels of cell density in the central medial (CM), reuniens (Re) and paraxiphoid (PaXi) nuclei, and medium labelling intensity in the CM, Re and paratenial (Pt) nuclei. In rats, Re and PaXi nuclei showed lower expression levels than in mice, although labeling intensity remained similar. In macaques, expression was generally high across all midline nuclei, with a slightly lower density in the Re and central latocellular (Clc) nuclei, and medium labelling intensity also lower in the central densocellular (Cdc) and central intermedial (Cim) nuclei (Fig 1 a1-c3; Fig 2 a1-c3).

### Intralaminar group

The intralaminar group showed the second highest FoxP2 expression across all species. Expression was strongest in the PF nucleus. In mice and rats, FoxP2 expression was higher in the centrolateral (CL) and parafascicular (PaF) nuclei than in the paracentral (PC) and angular (Ang) nuclei, with labeling intensity generally higher in rats than in mice. In the macaque, expression density was high in the central superior lateral (Csl) and parafascicular (Pf) nuclei, medium in the centromedian (CnMd) and lower levels in the paracentral (Pcn) and centrolateral (Cl) nuclei, while labeling intensity was the highest in the CnMd and Pf nuclei (Fig 1 a1-c3; Fig 2 a1-c3).

### Anterior group

The anterior group showed the largest differences in FoxP2 expression across species. In mice, FoxP2 was detected only in the interanterodorsal (IAD) and laterodorsal (LD) nuclei, with relatively few labeled cells and low labeling intensity. A few cells were observed in the anteromedial (AM) nucleus, but their number did not reach the threshold to be considered very low expression. In rats, expression increased across anterior nuclei, with cells detected in the anteroventral (AV) and AM nuclei, and higher density and labeling intensity in the IAD nucleus. In macaques, FoxP2 was expressed throughout all anterior thalamic nuclei, with low to medium cell density and labeling intensity. Overall, FoxP2 expressions in the anterior nuclei increased progressively from mice to rats to macaques. The most notable difference was observed in the LD nucleus, which had low cell density and labeling intensity in mice and rats, but medium-low density and medium intensity in macaques (Fig 1 a1-b3 ; Fig 2 a1-b3).

### Mediodorsal nucleus

FoxP2 expression in the mediodorsal (MD) nucleus was largely conserved across all species studied. Cell density was similar in mice, rats, and macaques, at a medium level, while labeling intensity was slightly higher in macaques, reaching medium-high levels. Within the nucleus, FoxP2 labeling intensity appeared higher in the medial MD than in the lateral MD (Fig 1 b1-c3; Fig 2 b1-c3).

### Ventral group

The ventral group showed notable differences in FoxP2 expression across species, with both cell density and labeling intensity increasing from mice to rats and then to macaques. In mice, FoxP2 density was very low across all ventral nuclei, with virtually no expression in the ventral posteromedial (VPM). Labeling intensity was generally low but slightly higher in the ventral anterior (VA), ventrolateral (VL), and submedius (Sub) nuclei compared to other ventral nuclei. In rats, FoxP2 density and intensity were higher than in mice, with the ventral posterior parvicellular component (VPPC) nucleus showing the highest expression. In macaques, cell density was similar to that in rats, but labeling intensity was higher in most ventral nuclei, except for the VA nucleus, which exhibited low cell density and intensity (Fig 1 a1-d3; Fig 2 a1-d3).

### Posterior group

FoxP2 expression in the posterior group was generally similar between mice and rats, with slightly higher labeling intensity in the suprageniculate (SG) nucleus of rats. In macaques, SG cell density was conserved, but labeling intensity was lower. The limitans (Li) nucleus in macaques exhibited higher cell density and labeling intensity than the posterior nuclei of rodents. FoxP2 expression in the lateral posterior (LP) nucleus was consistent across all species, with similar cell density and labeling intensity (Fig 1 c1-2, d1-e2; Fig 2 c1-2, d1-e2).

### Pulvinar group

FoxP2 is highly expressed in the pulvinar group in macaques, with the highest cell density observed in the pulvinar inferior (Pul I) and pulvinar oral (Pul O) nuclei, compared to lower density in the pulvinar medial (Pul M), pulvinar lateral (Pul L), and LP nuclei (Fig 3. e). Labeling intensity was relatively homogeneous across the pulvinar nuclei, with only a slightly lower intensity in Pul O (Fig 1 d3, e3; Fig 2 d3, e3).

**Fig 3.**
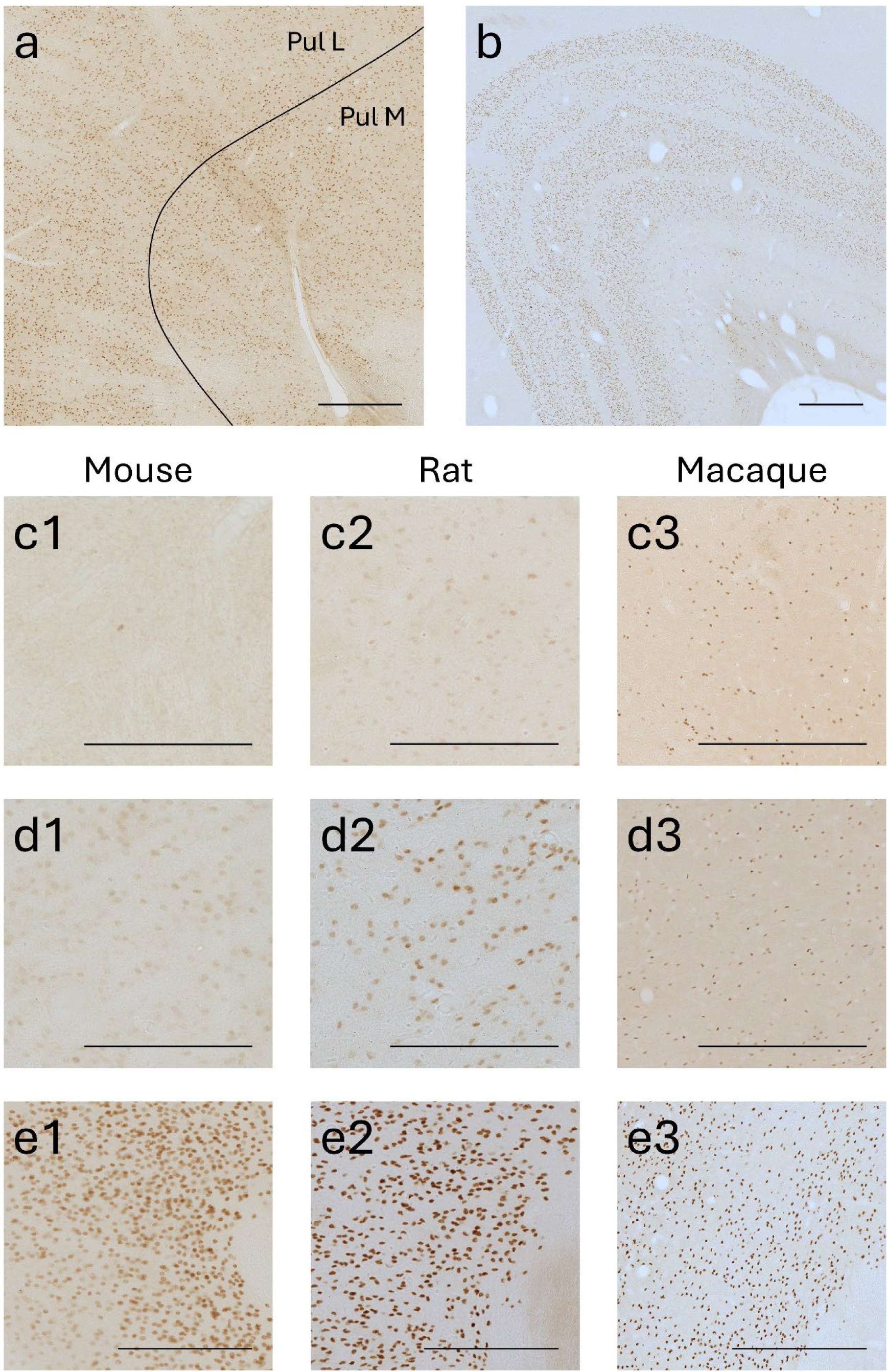
Representative images of FoxP2 labelling in the pulvinar (a) dorsal lateral geniculate (b) nuclei of macaque and the anteriomedial (c), ventral anterior (d) and parafascicular (e) nuclei of mouse (1), rat (2) or macaque (3). Scale bar: 500 μm

### Geniculate nuclei

FoxP2 expression in the medial geniculate (MG) nucleus was largely conserved across species, with similar cell density, although labeling intensity was higher in macaques and lower in rats. In contrast, expression in the dorsal lateral geniculate (DLG/GLd) nucleus showed pronounced species differences: mice and rats displayed relatively low FoxP2 expression, whereas macaques exhibited one of the highest levels. In macaques, FoxP2 expression was restricted to the parvocellular region, consistent with previous reports (Iwai et al., 2013) (Fig 3. d). Despite the high cell density in this region, likely influenced by cell size, labeling intensity remained relatively low. Expression in the macaque ventral lateral geniculate (GLv) nucleus was low with medium-low labelling intensity (Fig 1 c1-e2; Fig 2 c1-e2).

### Other

No FoxP2 expression was observed in R nucleus or the ZI in any of the studied species, nor in the pregeniculate (PrG) nucleus of mice. Rats showed low expression in the PreG nucleus. In mice, FoxP2 was absent in the lateral habenula (LH) but showed medium-low expression in the medial habenula (MH). In rats, expression was low in the lateral habenula and medium-high in the medial habenula. In both rodents, FoxP2-positive cells were concentrated in the lateral region of the MH. In macaques, both cell density and labeling intensity were low in the LH and MH (Fig 1 a1-e3; Fig 2 a1-e3).

In conclusion, FoxP2 is widely expressed in the thalamus of mice, rats and macaques with a wide range of FoxP2+ cell density and intensity. From almost no FoxP2 expression in nuclei like AM (Fig 3. a1-a3), medium expression levels in VA (Fig 3. B1-b3) and high expression in PaF (Fig 3. C1-c3).

## Discussion

In this study, we provide a comparative analysis of FoxP2 expression across thalamic nuclei in mice, rats and macaques. Expression was detected in most nuclei, with notable absence in the R nucleus and ZI, suggesting that FoxP2 is broadly involved in thalamic signaling but excluded from inhibitory regions. FoxP2 expression was particularly enriched in the midline and intralaminar nuclei, whereas the anterior thalamic group showed relatively weak labeling. Overall expression patterns were conserved between rodents and macaques, although several nuclei displayed species-specific differences. Macaques exhibited broader and, in some cases, more intense expression, in the anterior, ventral, pulvinar, and lateral geniculate nuclei, regions associated with higher-order sensory integration, except the lateral geniculate nuclei which is a relay nucleus.

FoxP2 expression was the highest in the *midline group* across all studied species, particularly in the PV, Rh and Xi nuclei of rodents and the PV, Cs and Cif nuclei of macaques. These nuclei project widely to the prefrontal and limbic cortex, hippocampus and amygdala forming a central component of the limbic-thalamic network involved in memory and learning, attention, behavioral flexibility and motivation (Joyce et al., 2022; Timbie et al., 2020; Vertes et al., 2015, 2022). Even though the midline nuclei have not directly been involved in language circuits, these processes are fundamental for supporting fluent language production and comprehension (Klostermann et al., 2013). The consistently high expression of FoxP2 across the studied species indicates a fundamental role in the formation and stabilization of thalamic circuits that support these integrative functions.

The *intralaminar group* showed the second highest levels of FoxP2 expression, with the highest expression in the PaF/Pf and CnMd nuclei. The striatum, hippocampus, and cerebellum are among the cortical and subcortical regions that these nuclei are closely linked to. Additionally, the PaF/Pf and CnMd project to visual, temporal, and frontal cortical regions (Eckert et al., 2011; Kumar et al., 2023). Thalamic aphasia has been linked to lesions in these nuclei, but there is evidence suggesting that these deficiencies may be caused by disturbances of pulvinar connection rather than damage to the thalamic nuclei themselves (Crosson, 2013). In addition, these nuclei are involved in arousal, emotion, cognition and motor processing, all of which may provide support for language functions (Kumar et al., 2023; Vertes et al., 2022). As in the midline nuclei, the elevated FoxP2 expression in the intralaminar nuclei points to a crucial role in establishing and maintaining their connectivity. Additionally, the variations in FoxP2 levels among individual midline and intralaminar nuclei may indicate functional specializations within these networks.

The *ventral group* showed notable differences across species, with FoxP2 levels increasing from rodents to macaques. The VA and VL have been found to be key structures in the cortico-striato-pallido-thalamo-frontal loops involved in motor speech production and fluency, as well as behavioral switching, timing and sequential processing (Barbas et al., 2013). Lesions or stimulation of these nuclei cause reduced speech output and fluency, perseverations and semantic paraphasias (Crosson, 2021a; Johnson & Ojemann, 2000). VA and VL have connections to the premotor and supplementary motor areas (Morel et al., 2005; Nakano et al., 1993). Additionally, VA is extensively connected to Brocás area (Bohsali et al., 2015). The increased expression of FoxP2 in macaques within these nuclei suggest further elaboration of these thalamo-cortical circuits, which may have allowed in humans for a more precise fine-tuning of motor sequencing and linguistic processing necessary for speech development and fluent communication.

FoxP2 expression in the *mediodorsal* nucleus was moderate across species, with slightly higher levels in the macaques. This nucleus projects extensively to the prefrontal cortex helping support behavioral flexibility, adaptive decision-making and memory (Mitchell, 2015; Vertes et al., 2015). Lesions to the MD are not commonly linked to aphasia, although they affect verbal memory, causing anomia, perseveration and executive deficits (Radanovic & Almeida, 2021), indicating an indirect contribution to linguistic organization and control. Despite this, Crosson proposed that the MD nucleus’s involvement in the cortico-striato-pallido-thalamo-frontal loops may contribute to linguistic procedural memory, which involves grammar (Crosson, 2021b). All in all, it seems that FoxP2 may be involved in supporting linguistic flexibility and adaptive control of speech by controlling synaptic efficiency and plasticity within the MD-prefrontal circuit.

The *pulvinar* nucleus displayed high FoxP2 expression in macaques. This structure, and especially its medial region, is involved in lexical-semantic integration for correct word selection, through its participation in cortico-thalamo-cortical circuits (Crosson, 2021a, 2021b). The Pul nucleus projects broadly to the cerebral cortex (Basile et al., 2024; Leh et al., 2008), including to Brocás area (Bohsali et al., 2015). Lesions to this region produce semantic paraphasias and fluent aphasia (Radanovic & Almeida, 2021), while stimulation can transiently interrupt naming (Ojemann, 1977). Thus, the pronounced FoxP2 expression in the macaque Pul indicates that FoxP2 likely contributes to the evolutionary precise regulation of cortico-thalamo-cortical feedforward and feedback loops that refine semantic representations.

FoxP2 expression was absent in the *reticular nucleus* and *zona incerta* across all species. The R nucleus regulates thalamic inhibition and attentional gating through GABAergic projections (Barbas et al., 2013; Crosson, 2013). The lack of Foxp2 expression in this nucleus implies that its role is not in inhibitory modulation, but rather in the relay and associative nuclei that maintain excitatory thalamocortical projections.

Taken together, these findings indicate that FoxP2 is preferentially expressed in thalamic nuclei that participate in key talamo-cortical circuits related to speech, memory and emotions. It is also relevant that FoxP2 is selectively highly expressed in the layer VI cortico-thalamic projection neurons (Hisaoka et al., 2010; Qi et al., 2024), which suggest that this transcription factor may be regulating both directions of the cortico-thalamo-cortical circuits, coordinating gene expression in neurons, allowing to maintain the precision of the reciprocal communication. Different studies support this hypothesis, as disruptions of FoxP2 expression on the cortex, striatum, cerebellum and thalamus affect the connectivity and plasticity of cortico-striatal, cerebellar, thalamo-striatal circuits (Chen et al., 2016; French et al., 2019; Rodríguez-Urgellés et al., 2023). Through its role as a transcription factor regulating neurogenesis, synaptic plasticity and axonal guidance (den Hoed et al., 2021), FoxP2 may be contributing to the fine-tuning of these circuits, ensuring the necessary temporal precision and cross-modal integration information across cortical, subcortical, and cerebellar components of speech-related networks.

FoxP2 expression studies performed in other mammals, such as echolocating bats (Rodenas-Cuadrado et al., 2018) confirm a great conservation of FoxP2 across the mammalian thalamic nuclei. No studies have been performed in the human adult thalamus, although, in the human fetal thalamus, FoxP2 has a similar expression pattern as in the macaque, with moderate-high Foxp2 expression in the pulvinar nucleus, high expression in the intralaminar, midline and ventral nuclei, medium expression in the MD and low to no expression in the anterior thalamic nuclei (Alhesain et al., 2025; Teramitsu et al., 2004).The most relevant finding of this studies is a high expression in the VA nucleus (Alhesain et al., 2025), which has a relatively low FoxP2 expression in the species studied in the present study. This reinforces the idea that during human evolution, FoxP2 expression has become selectively enhanced in thalamic territories associated with speech motor control and language processing. Given the role of the VA nucleus in the cortico-striato-pallido-thalamo-frontal loops (Barbas et al., 2013) and its dense connectivity with Brocás and premotor areas (Bohsali et al., 2015), this enhancement may reflect a molecular adaptation supporting the evolution of complex vocal behaviors and the modulation of motor-cognitive integration required for language.

One of the main differences between the primate (including human) thalamus and that of rodents is the selective expansion of associative nuclei, such as the pulvinar and mediodorsal nuclei, as well as the increased complexity and refinement of thalamo-cortical projections (Ruiz-Cabrera et al., 2023). These structural enlargements parallel the evolutionary expansion of association cortices and the emergence of higher-order cognitive and communicative abilities in primates. FoxP2 might have contributed to these evolutionary changes, as it is highly expressed in associative nuclei. Experimental studies in mice have shown that FoxP2 expression manipulation affects thalamic patterning. FoxP2 knock-out or R552H knock-in mutants exhibit a reduction of posterior thalamic regions and an expansion of intermediate nuclei (Ebisu et al., 2017), and similar reductions in thalamic volume have been observed in patients with FoxP2 mutations (Belton et al., 2003; Liégeois et al., 2016). Moreover, studies introducing humanized FoxP2 have found increased dendritic length of specifically thalamic, striatal and layer VI pyramidal neurons (Enard et al., 2009; Reimers-Kipping et al., 2011), highlighting its role in the evolutionary reorganization of cortico-basal ganglia circuits, in which the thalamus is involved.

Beyond evolution, these same molecular pathways may also confer vulnerability to neurodevelopmental and psychiatric disorders. Thalamocortical dysconnectivity is a common hallmark for multiple of these disorders, including schizophrenia bipolar disorder, major depressive disorders (MDD), autism spectrum disorder (ASD) and attention deficit and hyperactivity disorder (ADHD) (Giraldo-Chica & Woodward, 2017; Nair et al., 2015; Sheffield et al., 2020; Shi et al., 2025; Tomasi & Volkow, 2019; Tu et al., 2019). In these patients there is usually a reduction in thalamo-prefrontal connectivity and an increased thalamo-somatosensory connectivity. FoxP2 expression has been found to be reduced in schizophrenia and ASD patients, correlating with grey matter loss and cognitive and executive function deficits respectively (Haghighatfard et al., 2022; Sanjuán et al., 2021). Moreover, the polygenic risk score of FOXP2-related genes has been strongly associated with functional dysconnectivity identified in schizophrenia patients with shorter illness duration, which in term correlates with symptom score (Du et al., 2021). Our results show that FoxP2 expression is particularly enriched in associative nuclei which exhibit strong thalamocortical connectivity with prefrontal and association cortices, the same circuits that are seen to be disrupted in these disorders. A reduction or dysregulation of FoxP2 expression could alter the connectivity of these networks, leading to the language, executive and attentional impairments seen in these conditions. Thus, understanding FoxP2 expression in the thalamus not only informs us how language and cognitive abilities evolved but also how their dysfunctions can lead to disorders.

While these findings shed light on the potential role of FoxP2 in shaping thalamic circuits relevant to language and cognition, several limitations of this study should be considered. First, due to technical constraints we were unable to examine FoxP2 expression in adult human thalamic tissue. Current information is restricted to early human development (Alhesain et al., 2025; Teramitsu et al., 2004), although evidence from other species suggests that thalamic FoxP2 expression patterns established early in life are largely preserved into adulthood (Ferland et al., 2003; Takahashi et al., 2008). Second, the relationship between FoxP2 expression levels and establishment and function of thalamo-cortical circuits remains correlational. Further research combining manipulations of FoxP2 expression in vivo and axon-tracing would be necessary to determine how it translates into connectivity and circuit function. Finally, as language is a human-specific capacity, the specific contribution of individual thalamic nuclei to language circuits remains poorly defined, limiting our ability to infer the precise role of FoxP2 in these pathways.

In summary, FoxP2 is broadly expressed in the thalamus of rodents and primates, with conserved high expression in the midline and intralaminar nuclei and enhanced expression in associative thalamic nuclei in primates. This distribution mirrors the organization of language-related thalamocortical circuits, suggesting that FoxP2 may contribute to the molecular tuning of the circuits supporting language and speech.

## Author contributions

Tissue immunohistochemistry: BS-M, AU-H, MF-F. Image acquisition: BS-M, MF-F. Data analysis: BSM. Manuscript writing: BS-M, MAGC, CC, JG-J. All authors reviewed and approved the manuscript.

## Ethics declarations

The authors declare that they have no conflicts of interest.

The ethics protocol under which the experiments were approved was PROEX 209.7/22 and CEI-39-852. All animal experimentation was conducted in accordance with Directive 2010/63/EU of the European Parliament and of the Council of 22 September 2010 on the protection of animals used for scientific purposes and was approved by the Committee on Bioethics of the Universidad Autónoma de Madrid.

