## Supplementary Table 1 for "Comparative Analysis of FOXP2 Expression in the Thalamus of Mice, Rats, and Macaques: Implications for the Evolution of Language Circuits"

**Supplementary Table 1** Summary of FoxP2 cell density and intensity levels and number of analyzed sections for each thalamic nucleus in mice

| Thalamic nucleus | Density | | Intensity | | N |
|---|---|---|---|---|---|
| **Midline group** | | | | | |
| PV | 5 | | 5 | | 40 |
| IMD | 4 | | 4 | | 26 |
| IAM | 2 | | 1 | | 4 |
| CM | 3 | | 3 | | 40 |
| Rh | 5 | | 4 | | 28 |
| Re | 3 | | 3 | | 38 |
| PaXi | 3 | | 4 | | 28 |
| Xi | 5 | | 5 | | 28 |
| Pt | 4 | | 3 | | 6 |
| **Intralaminar group** | | | | | |
| Ang | 2 | | 2 | | 12 |
| CL | 4 | | 4 | | 34 |
| PC | 3 | | 3 | | 40 |
| PaF | 4 | | 5 | | 14 |
| **Anterior group** | | | | | |
| AD | 0 | | 0 | | 10 |
| AV | 0 | | 0 | | 12 |
| AM | 0 | | 1 | | 12 |
| IAD | 2 | | 1 | | 8 |
| LD | 1 | | 1 | | 18 |
| **Mediodorsal nucleus** | | | | | |
| MD | 3 | | 3 | | 38 |
| **Ventral group** | | | | | |
| VA | 1 | | 2 | | 6 |
| VL | 1 | | 2 | | 20 |
| VM | 1 | | 1 | | 34 |
| VPM | 0 | | 1 | | 41 |
| VPL | 1 | | 1 | | 42 |
| VPPC | 1 | | 1 | | 18 |
| Sub | 2 | | 2 | | 26 |
| **Posterior group** | | | | | |
| LP | 3 | | 3 | | 38 |
| Po | 2 | | 3 | | 51 |
| SG | 3 | | 3 | | 20 |
| **Geniculate nuclei** | | | | | |
| DLG | 1 | | 1 | | 37 |
| MG | 3 | | 3 | | 22 |
| **Other** | | | | | |
| Hm | 2 | | 2 | | 34 |
| Hl | 0 | | 2 | | 34 |
| PreG | 0 | | 0 | | 42 |
| R | 0 | | 0 | | 60 |
| ZI | 0 | | 0 | | 33 |

**Table 2** Summary of FoxP2 cell density and intensity levels and number of analyzed sections for each thalamic nucleus in rats

| Thalamic nucleus | Density | | Intensity | | N |
|---|---|---|---|---|---|
| **Midline group** | | | | | |
| PV | 5 | | 5 | | 38 |
| IMD | 4 | | 4 | | 22 |
| IAM | 2 | | 3 | | 2 |
| CM | 4 | | 4 | | 36 |
| Rh | 5 | | 5 | | 26 |
| Re | 2 | | 3 | | 36 |
| PaXi | 2 | | 3 | | 20 |
| Xi | 5 | | 4 | | 14 |
| Pt | 3 | | 4 | | 12 |
| **Intralaminar group** | | | | | |
| Ang | 2 | | 3 | | 8 |
| CL | 4 | | 4 | | 26 |
| PC | 3 | | 4 | | 34 |
| PaF | 5 | | 5 | | 10 |
| **Anterior group** | | | | | |
| AD | 0 | | 1 | | 14 |
| AV | 1 | | 1 | | 14 |
| AM | 1 | | 1 | | 14 |
| IAD | 3 | | 2 | | 4 |
| LD | 1 | | 1 | | 20 |
| **Mediodorsal nucleus** | | | | | |
| MD | 3 | | 3 | | 33 |
| **Ventral group** | | | | | |
| VA | 2 | | 2 | | 10 |
| VL | 2 | | 2 | | 18 |
| VM | 2 | | 2 | | 28 |
| VPM | 2 | | 2 | | 32 |
| VPL | 1 | | 2 | | 36 |
| VPPC | 3 | | 3 | | 12 |
| Sub | 2 | | 2 | | 24 |
| **Posterior group** | | | | | |
| LP | 3 | | 3 | | 34 |
| Po | 2 | | 3 | | 37 |
| SG | 3 | | 4 | | 6 |
| **Geniculate nuclei** | | | | | |
| DLG | 2 | | 1 | | 23 |
| MG | 3 | | 2 | | 14 |
| **Other** | | | | | |
| Hm | 4 | | 3 | | 23 |
| Hl | 1 | | 2 | | 19 |
| PreG | 1 | | 1 | | 24 |
| R | 0 | | 0 | | 42 |
| ZI | 0 | | 0 | | 40 |

**Table 3** Summary of FoxP2 cell density and intensity levels and number of analyzed sections for each thalamic nucleus in macaques

| Thalamic nucleus | Density | Intensity | N |
|---|---|---|---|
| **Midline group** | | | |
| Pa | 5 | 4 | 20 |
| Cs | 5 | 5 | 6 |
| Cdc | 4 | 3 | 10 |
| Cim | 4 | 3 | 9 |
| Cif | 4 | 5 | 9 |
| Clc | 3 | 3 | 10 |
| Ro | 4 | 4 | 5 |
| Re | 3 | 3 | 16 |
| Pt | 4 | 4 | 5 |
| **Intralaminar group** | | | |
| Csl | 4 | 3 | 9 |
| Pcn | 2 | 3 | 7 |
| Cl | 2 | 4 | 6 |
| CnMd | 3 | 5 | 11 |
| Pf | 4 | 5 | 5 |
| **Anterior group** | | | |
| AD | 1 | 1 | 11 |
| AV | 1 | 2 | 12 |
| AM | 1 | 2 | 9 |
| AI | 2 | 2 | 5 |
| LD | 2 | 3 | 9 |
| **Mediodorsal nucleus** | | | |
| MD | 3 | 4 | 9 |
| **Ventral group** | | | |
| VA | 1 | 1 | 10 |
| VL | 2 | 2 | 13 |
| VPL | 2 | 3 | 12 |
| VPM | 2 | 3 | 5 |
| VPMpc | 3 | 3 | 6 |
| VPI | 2 | 3 | 8 |
| X | 2 | 2 | 1 |
| **Posterior group** | | | |
| SG | 3 | 2 | 3 |
| Li | 4 | 4 | 1 |
| **Pulvinar group** | | | |
| Pul M | 3 | 3 | 3 |
| Pul L | 3 | 3 | 3 |
| Pul I | 4 | 3 | 4 |
| Pul O | 4 | 2 | 3 |
| LP | 3 | 3 | 5 |
| **Geniculate nuclei** | | | |
| GM | 3 | 4 | 5 |
| GLv | 1 | 2 | 3 |
| GLd | 5 | 1 | 11 |

| **Other** | | | | | |
|---|---|---|---|---|---|
| Hm | 1 | | 1 | | 2 |
| Hl | 1 | | 1 | | 2 |
| R | 0 | | 0 | | 25 |
| ZI | 0 | | 0 | | 9 |
